# TASOR epigenetic repressor cooperates with a CNOT1 RNA degradation pathway to repress HIV

**DOI:** 10.1101/2020.11.22.386656

**Authors:** Roy Matkovic, Marina Morel, Pauline Larrous, Benjamin Martin, Fabienne Bejjani, Stéphane Emiliani, Sarah Gallois-Montbrun, Florence Margottin-Goguet

**Affiliations:** Inserm, U1016, Institut Cochin, 22 rue Méchain, 75014 Paris, France; CNRS, UMR8104, Paris, France; Université de Paris, Paris, France

## Abstract

The Human Silencing Hub (HUSH) complex constituted of TASOR, MPP8 and Periphilin is involved in the spreading of H3K9me3 repressive marks across genes and transgenes such as ZNF encoding genes, ribosomal DNAs, LINE-1, Retrotransposons and Retroelements or the integrated HIV provirus^1–5^. The deposit of these repressive marks leads to heterochromatin formation and inhibits gene expression. The precise mechanisms of silencing mediated by HUSH is still poorly understood. Here, we show that TASOR depletion increases the accumulation of transcripts derived from the HIV-1 LTR promoter at a post-transcriptional level. By counteracting HUSH, Vpx from HIV-2 mimics TASOR depletion. With the use of a Yeast-Two-Hybrid screen, we identified new TASOR partners involved in RNA metabolism including the RNA deadenylase CCR4-NOT complex scaffold CNOT1. TASOR and CNOT1 interact *in vivo* and synergistically repress HIV expression from its LTR. In fission yeast, the RNA-induced transcriptional silencing (RITS) complex presents structural homology with HUSH. During transcription elongation by RNA polymerase II, RITS recruits a TRAMP-like RNA degradation complex composed of CNOT1 partners, MTR4 and the exosome, to ultimately repress gene expression *via* H3K9me3 deposit. Similarly, we show that TASOR interacts and cooperates with MTR4 and the exosome, in addition to CNOT1. We also highlight an interaction between TASOR and RNA Polymerase II, predominantly under its elongating state, and between TASOR and some HUSH-targeted nascent transcripts. Furthermore, we show that TASOR overexpression facilitates the association of the aforementioned RNA degradation proteins with RNA polymerase II. Altogether, we propose that HUSH operates at the transcriptional and post-transcriptional levels to repress HIV proviral gene expression.

---

As of today, highly active antiretroviral therapy (HAART) is very efficient in inhibiting the replication of HIV in CD4^+^ infected T cells thanks to the combination of several molecules targeting different stages of the lentivirus cycle. As long as the treatment is properly followed, the viremia of the infected individual will become undetectable. However, if treatment is randomly taken or is interrupted, a viral rebound will cause new cellular infections’ increasing the number of reservoir cells in which the virus remains latent. At the proviral level, latency can be explained by an inhibition of viral transcription or by a defect in the post-transcriptional steps leading to a lack of production of new lentiviral particles. After integration, HIV-1 transcription involves cellular class II transcriptional complexes, as well as HIV proteins such as Transactivator of transcription Tat protein, to ensure the production of genomic RNA and splice variants leading to viral production. However, HIV-1 integration in poorly transcribed, desert, geneless, or centromeric regions can result in the repression of HIV-1 expression due to multiple epigenetic mechanisms^6–9^. The HUSH complex-TASOR, MPP8 and periphilin-was identified as a potential player in HIV repression, propagating repressive H3K9 repressive marks in a position-effect variegation manner in a HIV-1 model of latency^1,3^. We and others have previously shown that Vpx of SIVsmm/HIV-2 could counteract HUSH by preferentially degrading TASOR and MPP8^3,10^. This degradation leads to a decrease in the HUSH dependent-H3K9me3 repressive marks on the coding sequence, and then to an increase in the amount of RNA transcribed from the LTR promoter of the virus^3^. This increase was improved under TNFα-treatment, a well-known transcriptional activator, which led us to question a possible co-or post-transcriptional repressive effect of TASOR. Another clue towards this hypothesis was the discovery of the interaction of TASOR (*alias* C30rf63) and Periphilin-1 with messenger RNAs^11,12^, or with the *XIST* lncRNA^13^.

To address the question of a possible co- or post-transcriptional repressive effect of TASOR toward HIV, we chose a cellular system in which the LTR-driven active transcription could be studied independently of RNA splicing. Therefore, we used HeLa cells harboring one unique and monoclonal copy of an integrated LTR-Luciferase construct with a deleted TAR sequence (ΔTAR) to get a fully active transcription process^14^ (Fig. 1a). Indeed, due to the removal of the TAR RNA structure that blocks the elongating process of RNA polymerase II (RNAPII), these cells express around a 100 time more Luciferase (Luc) transcripts than the WT LTR cells in the absence of Tat (Fig. 1a). Consequently, we showed that TASOR still had a repressive role in these cells by measuring the Luc activity when we overexpressed TASOR, or in contrast, when we inhibited its expression by siRNA (Fig. 1b, 1c). To better characterize this repression, we conducted *Nuclear Run On* experiments to examine the role of TASOR on the mRNA metabolism process. This method enables to separate the repressive role of TASOR at the transcriptional level from its potential post-transcriptional role, by comparing the BrUTP-incorporated nascent Luc transcripts to the total amounts of Luc mRNA (Fig. S1a). While TASOR downregulation by siRNA very slightly increases by a 1.3-fold the nascent transcripts (Fig. 1d), as also seen on WT LTR cells (Fig. S1b), it triggers a 2.6-fold accumulation of total RNA in the cells, suggesting that TASOR depletion might impair the turnover of these LTR-driven transcripts (Fig. 1d). Mimicking TASOR depletion, Vpx WT from HIV-2, which depletes TASOR (Fig. 1e, left), increases Luc RNA synthesis by inducing a 2-fold more accumulation of the LTR-driven transcript at a post-transcriptional step (2.4-fold effect on the nascent and 5.5 on the total), again suggesting an impact on transcripts stability (Fig. 1e, S1c, S1d). However, Vpx WT solely increased the transcription of a cellular mRNA such as the MORC2 RNA, a known cellular target of HUSH^15^ (Fig. S1d, middle). These effects on Luc or MORC2 RNA were lost with a Vpx mutant unable to induce TASOR degradation (Fig. S1c and S1d). We have also performed the experiment with Vpr from HIV-1, which presents some structural similarities with Vpx, while unable to induce TASOR depletion^16,3^. Vpr was not able to induce TASOR degradation as expected, nor to increase the expression of the transcript at any stage in this LTR-TAR-deleted system but, in contrast to Vpx, it specifically stabilized the TNFα transcripts, in agreement with the previously described effect of Vpr on TNFα production^17^ (Fig. S1c and S1d, right). These results suggest that Vpx is able to stabilize LTR-driven transcripts *via* TASOR degradation. Structure/function prediction analyses of TASOR first 900 amino acid (aa) sequence using the PSIPRED server (UCL Bioinformatics group) revealed a PARP13-like PARP domain at its N-terminus region (300 aa) with high probable and reliable functions in mRNA binding, splicing and processing, together with the SPOC (Spen paralog and ortholog C-terminal) domain, often associated with transcription repression (Table 1). The bioinformatic structure prediction and amino-acid alignment of different SPOC domains show that TASOR’s SPOC domain is homologous to the one found in *Arabidopsis thaliana* FPA protein, a protein described to be involved in epigenetic repression of retrotransposons and to suppress intergenic transcription in an RNA-dependent manner^18,19^ (Fig. S2a, S2b). To get more details on how TASOR could inhibit the turnover of the LTR-driven Luc transcripts, we performed a Yeast-Two-Hybrid (Y-2-H) screen using as a bait the first 900 aa of the 1670 aa-long TASOR protein, and as a prey the proteome from human macrophages which are natural HIV target cells. The results show that TASOR N-terminus region binds its known HUSH partner, MPP8. Other candidate binding-proteins are factors involved in gene transcription modulation, LXRα, ARID1A known as BAF250, EP300, SUPT6H, KDM5C, and proteins involved in mRNA metabolism, CYFIP1, CNOT1, DHX29, DHX30, EIF4G1, MATR3 (Fig. S2c). The latter has already been discovered as a TASOR interacting protein in large scale interactome studies^20,21^. Among these candidate partners of TASOR, some are negative regulatory factors of gene expression directly connected to RNA pol II (RNAPII) subunit RPB1 (Fig. 2a), which may indicate a role of TASOR and partners in inhibiting gene expression during the transcription process *per se*. We chose to focus on CNOT1, which is the scaffold protein of the most important and conserved deadenylase complex from Yeast to Human, the Carbon catabolite repression 4-negative on TATA-less complex or CCR4-NOT^22,23^. We confirmed the interaction between TASOR-DDK with endogenous CNOT1, besides already known TASOR-binding proteins such as MPP8, periphilin and MORC2 (Fig. 2b). While DNase I did not inhibit TASOR and CNOT1 interaction, RNase A did decrease their interaction in HeLa cells harboring or not the LTR-ΔTAR-Luc transgene (Fig. 2c and Fig. S2d), suggesting that the presence of an RNA favors the interaction between the two proteins. CNOT1 being a scaffold protein, we wanted to check whether TASOR could interact with another component of the CCR4-NOT complex. Because we could not efficiently overexpress CNOT1, we chose the deadenylase CNOT7 partner to pull-down the complex in the reverse situation. We immunoprecipitated the DDK-tagged CNOT7 and revealed an interaction with endogenous TASOR, and the other CCR4-NOT complex components CNOT9 and CNOT1 proteins (Fig. 2d).

**Figure 1:**
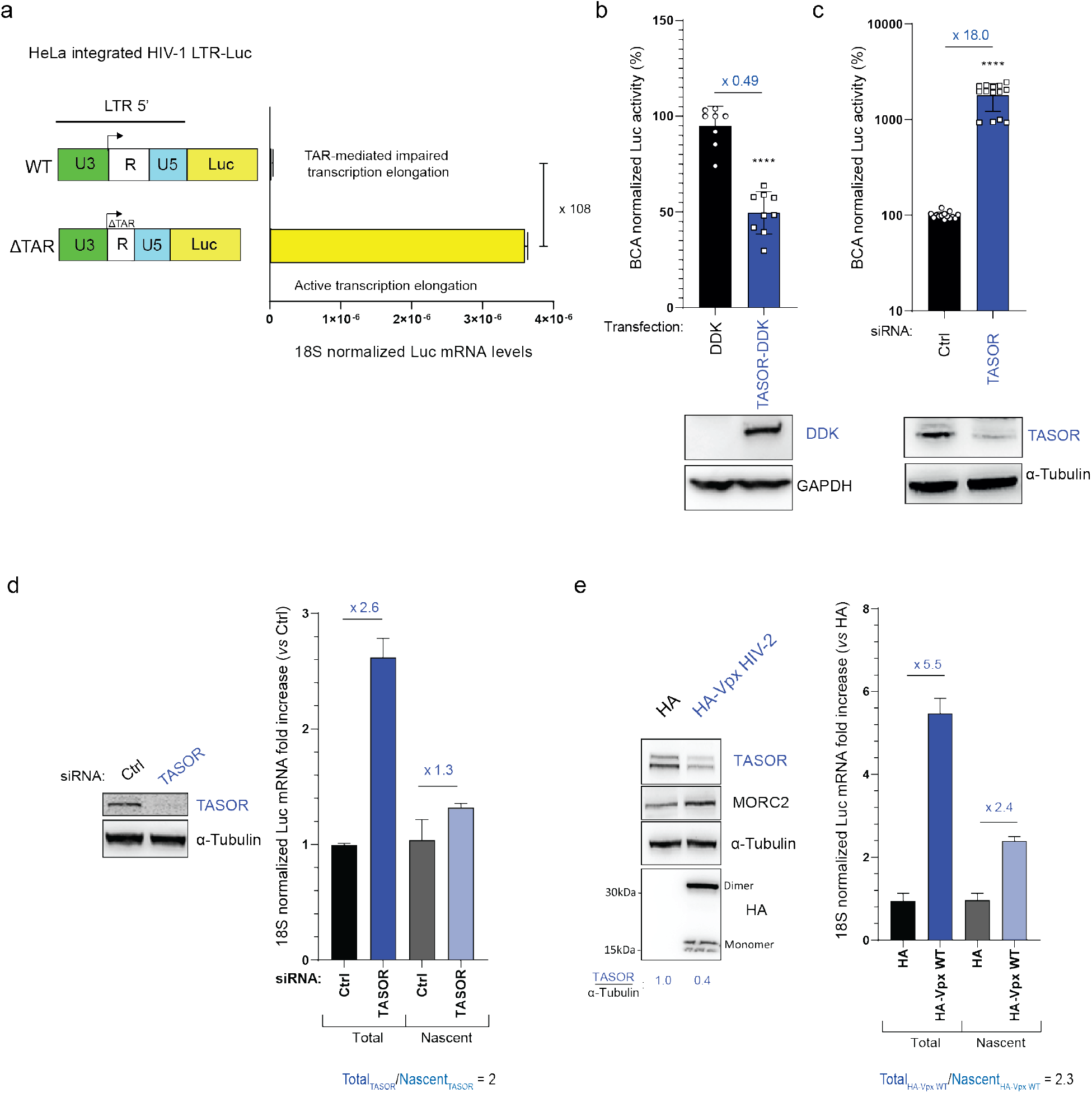
TASOR destabilizes HIV-1 LTR-driven transcripts. **a** HeLa HIV-1 LTR-ΔTAR-Luc cell model. HeLa HIV-1 LTR-Luc and HeLa HIV-1 LTRΔTAR-Luc cells were lysed for Luc RNA quantification. **b** TASOR over-expression decreases LTR-driven Luc expression. HeLa HIV-1 LTRΔTAR-Luc were lysed 48h after pLenti-DDK or pLenti-TASOR-DDK transfection and Luc activity was measured. Two-sided unpaired t test was performed (n = 3, ****p < 0.0001). **c** siRNA-mediated TASOR depletion increases LTR-driven Luc expression. HeLa HIV-1 LTRΔTAR-Luc were lysed 48h after siCtrl or siTASOR transfection and Luc activity was measured. Two-sided unpaired t test was performed (n = 3, ****p < 0.0001). **d** TASOR negatively impacts LTR-driven Luc transcript at a post-transcriptional step. *Nuclear Run On* was performed in HeLa HIV-1 LTRΔTAR-Luc after 72h of siCtrl or siTASOR transfection. Data show a representative experiment (from three independent experiments-see Fig. 2e for more data). **e** HIV-2 Vpx mimics siRNA-mediated TASOR depletion in increasing LTR-driven transcript stability. Nuclear Run On performed in HeLa HIV-1 LTRΔTAR-Luc after 48h of pAS1B-HA or pAS1B-HA-Vpx Ghana transfection. Data show one representative experiment (from three independent experiments-see Fig. S1d for more data).

**Figure 2:**
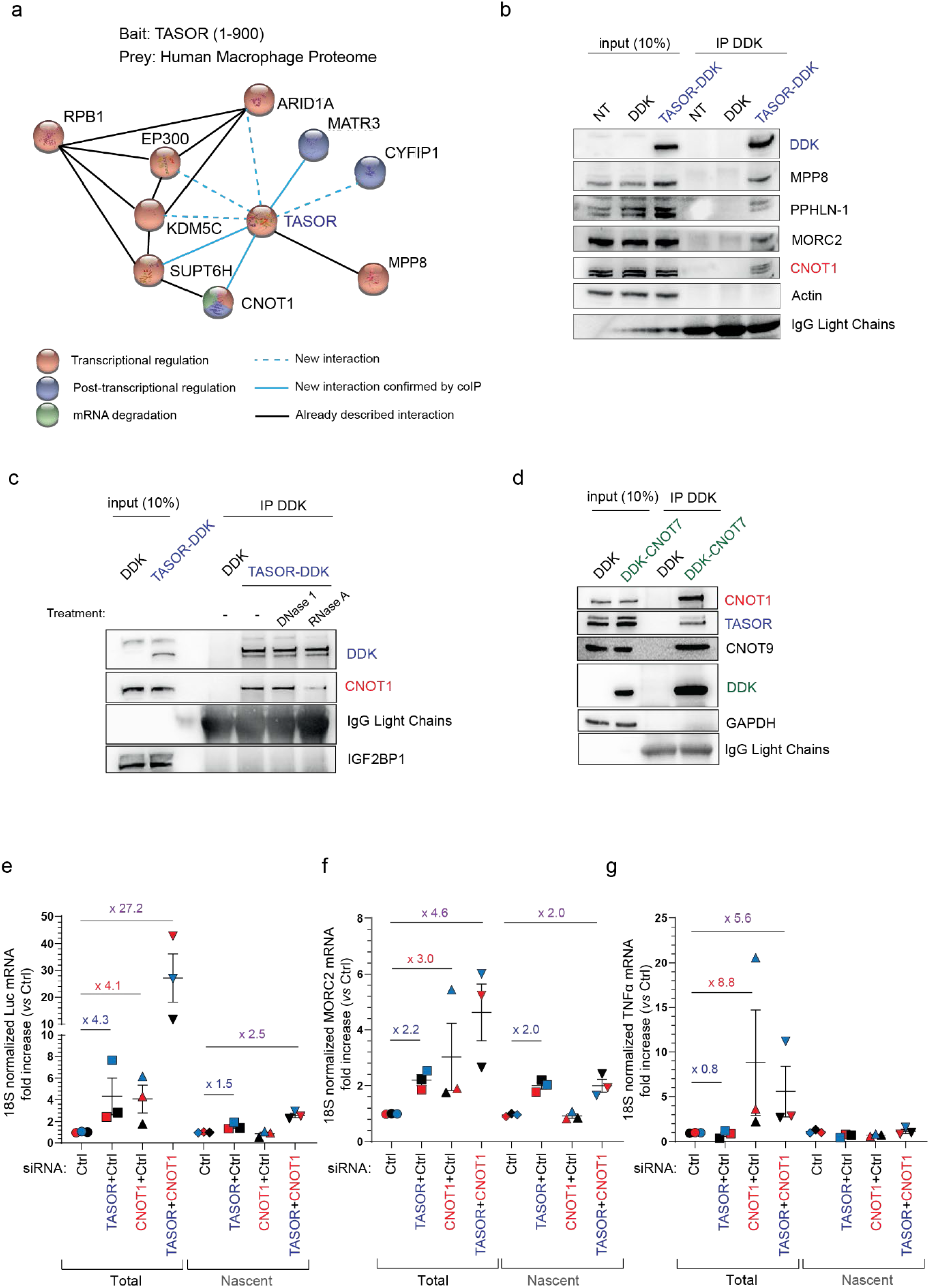
TASOR interacts and cooperates with the CCR4-NOT complex scaffold CNOT1 to destabilize LTR-driven transcripts. **a** Identification of CNOT1 and other proteins as new partners of TASOR by Yeast-two-Hybrid screening. Network of interactions was built using STRING. Proteins involved in transcriptional regulation, in post-transcriptional regulation of RNAs, and in mRNA degradation pathways are colored in red, blue and green respectively. Newly found interactions are shown in blue. Those validated by coIP in this manuscript are shown in solid line (TASOR-CNOT1 validated in Fig. 2b, 2c, 3b, 2c, TASOR-SUPTH6 in Fig. S2g and TASOR-Matrin 3 in Fig. 3c, 4g). **b** TASOR interacts with CNOT1 by co-immunoprecipitation. Mock, DDK, TASOR-DDK vectors were transfected in HeLa cells and anti-DDK immunoprecipitation was performed. **c**, A transcript stabilizes TASOR-CNOT1 interaction. DDK, TASOR-DDK vectors were transfected in HeLa cells. Lysates were treated or not with DNase 1, or RNase A. Anti-DDK immunoprecipitation was then performed. IGF2BP1 is a negative control. **d** TASOR interacts with the CCR4-NOT complex deadenylase CNOT7. DDK and TASOR-DDK vectors were transfected in HeLa cells and anti-DDK immunoprecipitation was performed. **e** TASOR and CNOT1 cooperate to repress HIV-1 LTR expression at a post-transcriptional level. After siRNA transfection in HeLa HIV-1 LTR-ΔTAR-Luc, a *Nuclear Run On* experiment was undertaken to measure LTR-driven Luc transcripts levels. (n=3; each color represents one different independent experiment, mean and SEM are showed). **f** TASOR represses MORC2 transcription while CNOT1 decreases its stability. After siRNA transfection in HeLa HIV-1 LTR-ΔTAR-Luc, a *Nuclear Run On* experiment was undertaken to measure MORC2 transcripts levels. (n=3; each color represents one different independent experiment, mean and SEM are showed). **g** CNOT1 depletion only increases TNFα transcript stability. After siRNA transfection in HeLa HIV-1 LTR-ΔTAR-Luc, a *Nuclear Run On* experiment was undertaken to measure TNFα transcripts levels. (n=3; each color represents one different independent experiment, mean and SEM are showed).

**Table 1:** Domain fold recognition and eukaryotic function predictions of TASOR 1-900. The first 900 amino-acid sequence of TASOR (NCBI Reference Sequence: NP_001106207.1) was loaded on the PSIPRED server (http://bioinf.cs.ucl.ac.uk/psipred/) for Domain fold recognition (pGenTHREADER) and Eukaryotic function predictions (FFPred3). Results were sorted according to confidence/reliability.

| pGenTHREADER, Fold Recognition | TASOR 1-900 aa |  |  |
| --- | --- | --- | --- |
| Confidence | Target protein (Human unless specified) | Net Score | p-value |
| CERT | PARP12 | 68.105 | 7,00E-06 |
| CERT | PARP14 | 67.410 | 8,00E-06 |
| CERT | PARP15 | 61.019 | 3,00E-05 |
| CERT | PARP13 | 58.639 | 6,00E-05 |
| HIGH | PARP1 | 52.145 | 3,00E-04 |
| HIGH | PARP10 | 47.367 | 8,00E-04 |
| MEDIUM | RCD1 (Arabidopsis Thaliana) | 45.574 | 1,00E-03 |
| MEDIUM | RNA Helicase Aquarius | 44.130 | 2,00E-03 |
| MEDIUM | ADRM1 | 43.744 | 2,00E-03 |
| MEDIUM | Spoc domain of FPA (Arabidopsis Thaliana) | 43.610 | 2,00E-03 |
| MEDIUM | Ovotransferrin (Anas platyrhynchos) | 43.475 | 2,00E-03 |
| MEDIUM | Anthrax Edema Factor | 43.414 | 2,00E-03 |
| MEDIUM | Ovotransferrin (Gallus gallus) | 40.177 | 4,00E-03 |
| MEDIUM | Ku | 39.673 | 5,00E-03 |
| MEDIUM | C-Src | 39.475 | 5,00E-03 |

**Table 1: Domain fold recognition and eukaryotic function predictions of TASOR 1-900.**
| Biological Process Predictions | TASOR 1-900 aa |  |  |
| --- | --- | --- | --- |
| GO term | Biological process | Prob | SVM reliability |
| GO:0008380 | RNA splicing | 0.937 | High |
| GO:0019222 | regulation of metabolic process | 0.918 | High |
| GO:0000398 | mRNA splicing, via spliceosome | 0.891 | High |
| GO:0000375 | RNA splicing, via transesterification reactions | 0.868 | High |
| GO:0034645 | cellular macromolecule biosynthetic process | 0.832 | High |
| GO:0006396 | RNA processing | 0.813 | High |
| GO:0006351 | transcription, DNA-templated | 0.813 | High |
| GO:2001141 | regulation of RNA biosynthetic process | 0.749 | High |
| GO:1903506 | regulation of nucleic acid-templated transcription | 0.740 | High |
| GO:0006397 | mRNA processing | 0.734 | High |
| GO:0010468 | regulation of gene expression | 0.676 | High |
| GO:0051171 | regulation of nitrogen compound metabolic process | 0.633 | High |
| GO:0006810 | transport | 0.614 | High |
| GO:0010629 | negative regulation of gene expression | 0.578 | High |
| GO:0009059 | macromolecule biosynthetic process | 0.568 | High |
| GO:0051252 | regulation of RNA metabolic process | 0.527 | High |
| GO:0006355 | regulation of transcription, DNA-templated | 0.517 | High |
| GO:0044237 | cellular metabolic process | 0.954 | Low |
| GO:0008152 | metabolic process | 0.922 | Low |
| GO:0006807 | nitrogen compound metabolic process | 0.869 | Low |
| GO:0034641 | cellular nitrogen compound metabolic process | 0.868 | Low |
| GO:0006139 | nucleobase-containing compound metabolic process | 0.868 | Low |
| GO:0046483 | heterocycle metabolic process | 0.838 | Low |
| GO:0006725 | cellular aromatic compound metabolic process | 0.833 | Low |
| GO:0009058 | biosynthetic process | 0.815 | Low |

Since TASOR and CNOT1 interact with each other, we wondered whether they both collaborated to repress the LTR-driven transcripts expression at the level of the transcription *per* se and/or at the level of RNA stability by performing *Nuclear Run On* experiments as in Fig. 1. Depletion of TASOR and/or CNOT1 was checked by western-blot (Fig. S2e). As previously observed for TASOR, the major effect of CNOT1 was detected at the total level of LTR-driven transcripts, which were increased upon CNOT1 depletion, in agreement with its known role on RNA stability (Fig. 2e). Interestingly, when inhibiting both proteins expression, a strong synergistic effect was observed at the stability level, suggesting that TASOR and CNOT1 act in different pathways but cooperate to repress the expression of the transcript particularly at a post-transcriptional level (Fig. 2e: 27.2-fold on total RNA and 2.5-fold on nascent RNA). Importantly, siRNA/Vpx-mediated TASOR depletion only increased the level of MORC2 RNAs at the transcriptional stage (no difference between total and nascent RNA), in agreement with previously published data showing an epigenetic silencing of MORC2 gene by the HUSH complex^15^ (Fig. 2f; Fig. S1d, middle). In contrast, CNOT1 depletion did not increase nascent MORC2 mRNA levels but led to its accumulation (Fig. 2f). On the other hand, CNOT1 has been shown to interact with the TNFα mRNA and to induce its deadenylation with CNOT7^24,25,26,27^. Accordingly, we then confirmed that CNOT1 depletion had an effect at the total RNA level without influencing TNFα transcription (Fig. 2g). Of note, the combination of siCNOT7 and siCNOT1 did not better increase Luc expression than the siCNOT1 alone, in agreement with CNOT1 and CNOT7 belonging to the same pathway (Fig. S2f). However, the inhibition of both CNOT7 and TASOR expression confirmed the collaboration between CCR4-CNOT and TASOR (Fig. S2f). Altogether, our results show that TASOR and CCR4-NOT interact both physically and functionally and can then strongly cooperate at a post-transcriptional level to decrease LTR-driven transcript accumulation.

To gain insight into the mechanism of TASOR-CNOT1-mediated repression mechanism, we questioned the involvement of MTR4 from the TRAMP/NEXT complex, Exosc10/RRP6 from the exosome and the m^6^A reader YTHDF2. Indeed, in yeast, Not1, the human CNOT1 counterpart, has been shown to recruit MTR4 and the exosome to induce the degradation of nuclear defective and nascent RNAs^28,29^. In addition, MTR4 and Exosc10/RRP6 are members of an RNA surveillance complex that inhibits LTR-driven expression in the HeLa HIV-1 WT LTR-Luc model^30^. Finally, YTHDF2 was reported on the one hand to interact with CNOT1, which leads to the destabilization of m^6^A modified mRNAs^31^ and on the other hand, to be recruited onto HIV-1 5’LTR, Nef and 3’LTR genomic RNA sequences^32,33^. By cell fractionation, we showed that TASOR was mainly found in the nucleus of human cells, alike the nuclear MATR3 protein, while CNOT1 is a shuttling protein, present both in the cytoplasm, alike GAPDH, and in the nucleus (Fig. 3a). Then, in an endogenous CNOT1 immunoprecipitate from nuclear extracts, we could retrieve CNOT7 and YTHDF2 as expected, as well as MTR4, Exosc10, TASOR (Fig. 3b). The same components were pulled-down in a MTR4 immunoprecipitate except CNOT7 (Fig. 3b). Next, we overexpressed TASOR-DDK in cells and performed a DDK immunoprecipitation to recover TASOR and its bound partners in the absence/presence of RNase A. MATR3 is the only protein along with MPP8, our positive control, to be efficiently co-immunoprecipitated with TASOR under treatment with RNase A, which confirms our Y-2-H results and that this interaction is RNA-independent (Fig. 3c). TASOR interacts with the CNOT1 partners MTR4, YTHDF2 and, as for CNOT1, these interactions are destabilized in the presence of RNase A (Fig. 3c). TASOR also interacts with Exosc10 (Fig. S3a)

**Figure 3:**
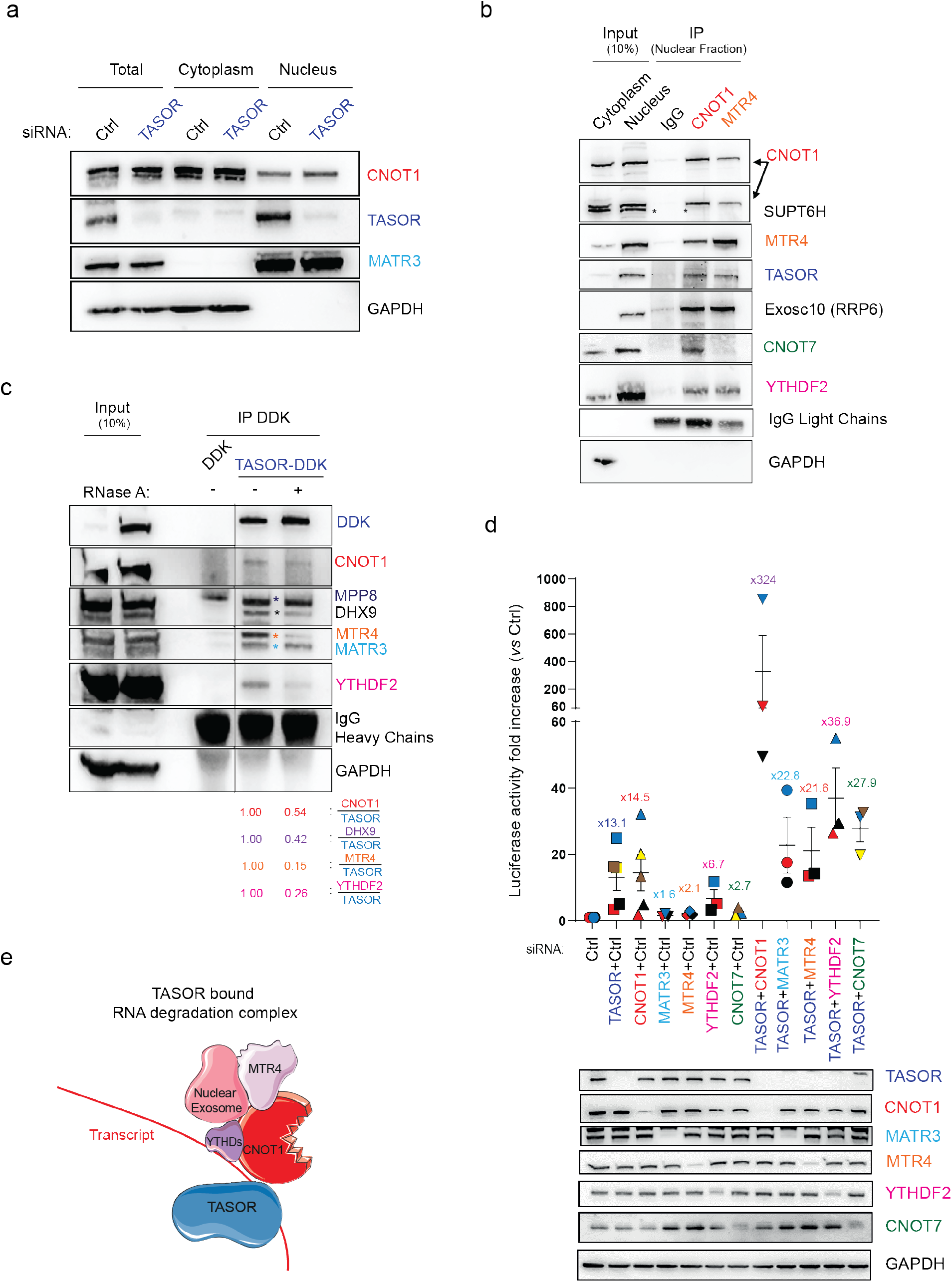
TASOR interacts and cooperates with nuclear RNA degradation factors. **a** TASOR is a nuclear protein. Cytoplasmic and nuclear protein extracts from HeLa HIV-1 LTR-ΔTAR-Luc cells were loaded on a SDS-PAGE gel. GAPDH and MATR3 are markers of the cytoplasmic and nuclear fractions respectively. **b** TASOR interacts nuclear CNOT1 and the TRAMP-like/NEXT component MTR4. HeLa HIV-1 LTR-ΔTAR-Luc cells were fractionated and endogenous CNOT1, MTR4 immunoprecipitation was performed in the nuclear fraction. The asterisk shows the SUPT6H band. **c** TASOR interacts with the known CNOT1 partners in an RNA-dependent manner. DDK, TASOR-DDK vectors were transfected in HeLa HIV-1 LTR-ΔTAR-Luc cells. After 48 h, lysates were treated or not with RNase A for 30min at Room Temperature. Anti-DDK immunoprecipitation was then performed. All lanes are from the same gel. **d** TASOR cooperates with the nuclear RNA destabilization/degradation factors. After 72h of siRNA transfections in HeLa HIV-1 LTR-ΔTAR-Luc, cells were lysed and Luciferase activity was measured and normalized on protein concentration (n=5 for the siRNA TASOR and siRNA CNOT1 conditions; n=3 for other siRNA transfections; each color represents one different independent experiment, mean and SEM are showed). **e** Schematic representation of RNA-mediated interactions between TASOR and CNOT1 and its partners: the TRAMP-like/NEXT components MTR4 and Nuclear Exosome, and the m^6^A reader YTHDF2.

To verify the involvement of these new factors in the TASOR repression of LTR expression, we depleted these proteins alone or in a combination with TASOR and measured the Luciferase expression (Fig. 3d). The strongest synergistic effect was always obtained under TASOR and CNOT1 co-depletion (Fig. 3d). Nonetheless, we also observed a synergistic repression effect among all the proteins tested, namely between TASOR and MATR3, MTR4 or YTHDF2 (Fig. 3d) or Exosc10 (Fig. S3b). In conclusion, our results suggest that TASOR and CNOT1 together form a platform recruiting different factors involved in RNA degradation pathways (Fig. 3e).

The ability of TASOR to propagate H3K9me3 marks and its association with CNOT1 and subunits of a TRAMP-like/NEXT complex and the exosome led us to imagine HUSH as a sort of human counterpart of the fission yeast RNA-induced silencing complex (or RITS). Indeed, RITS induces transcriptional silencing by addition of H3K9me3 marks concomitantly to the synthesis of a transcript and its processing by a TRAMP complex and the exosome^34^. Alternatively, silencing depends on recognition of the transcript by a small interfering RNA produced by the fission yeast dcr1 dsRNA-cleavage activity^35,36^. The inhibition of DICER by siRNAs did not increase the LTR-driven Luc expression (Fig. S3c) suggesting that the RNAi pathway is not involved in our system. In addition, RITS is constituted of Chp1 and Tas3, with Chp1 harboring a SPOC domain alike TASOR and a trimethyl-binding domain, the chromodomain, alike MPP8^37,38^. Therefore, we hypothesized that HUSH could propagate epigenetic marks by following the RNA polymerase II. The analysis of ChIP-seq and RNA-seq data from Liu *et al*^2^ supported this hypothesis, as we could find several examples of genes covered by RNAPII and TASOR-dependent H3K9me3 marks (Fig. 4a, Fig. S4a, S4b). Next, we could recover RNAPII in a TASOR-immunoprecipitate (Fig. 4b). More importantly, we uncover that TASOR interacted predominantly with a phosphorylated form of RNAPII, PhosphoSer2-RNAPII, specific to the elongation phase of transcription (Fig. 4c, Fig. S4c), rather than with the phosphorylated form enriched during transcription initiation, PhosphoSer5-RNAPII (Fig. 4c, S4c). Next, we used Formaldehyde Assisted Isolation of Regulatory Elements coupled with qPCR (FAIRE-qPCR), which allows to distinguish nucleosome-depleted DNA regions from chromatin. While TASOR depletion had no impact on the so-called Nuc0 nucleosome positioned at the very beginning of the LTR promoter, it increased chromatin accessibility of the LTR-Nuc1 region downstream of the Transcription start site (TSS) and triggered a dramatic decompaction of the Luc coding sequence, even more visible than the change induced by TNFα itself (Fig. 4d and Fig. 4e). Strikingly, overexpression of TASOR enhanced the binding of CNOT1, MTR4, Exosc10, YTHDF2 and MORC2 on RNAPII but not that of the RNAPII elongation cofactor SUPT6H (Fig. 4f). Furthermore, following BrUTP labeling of nascent transcripts, we coupled immunoprecipitation of TASOR-DDK with RT-qPCR and found that the nascent Luc transcript, derived from the LTR, the nascent MORC2 transcript and the nascent TUG1 lncRNA, which we identified as a TASOR’s target with the data from Liu *et al*. (Fig. 4a), were associated with TASOR (Fig. 4g). Altogether, these results suggest that TASOR helps assembling RNA metabolism machineries on RNAPII during transcription elongation.

**Figure 4:**
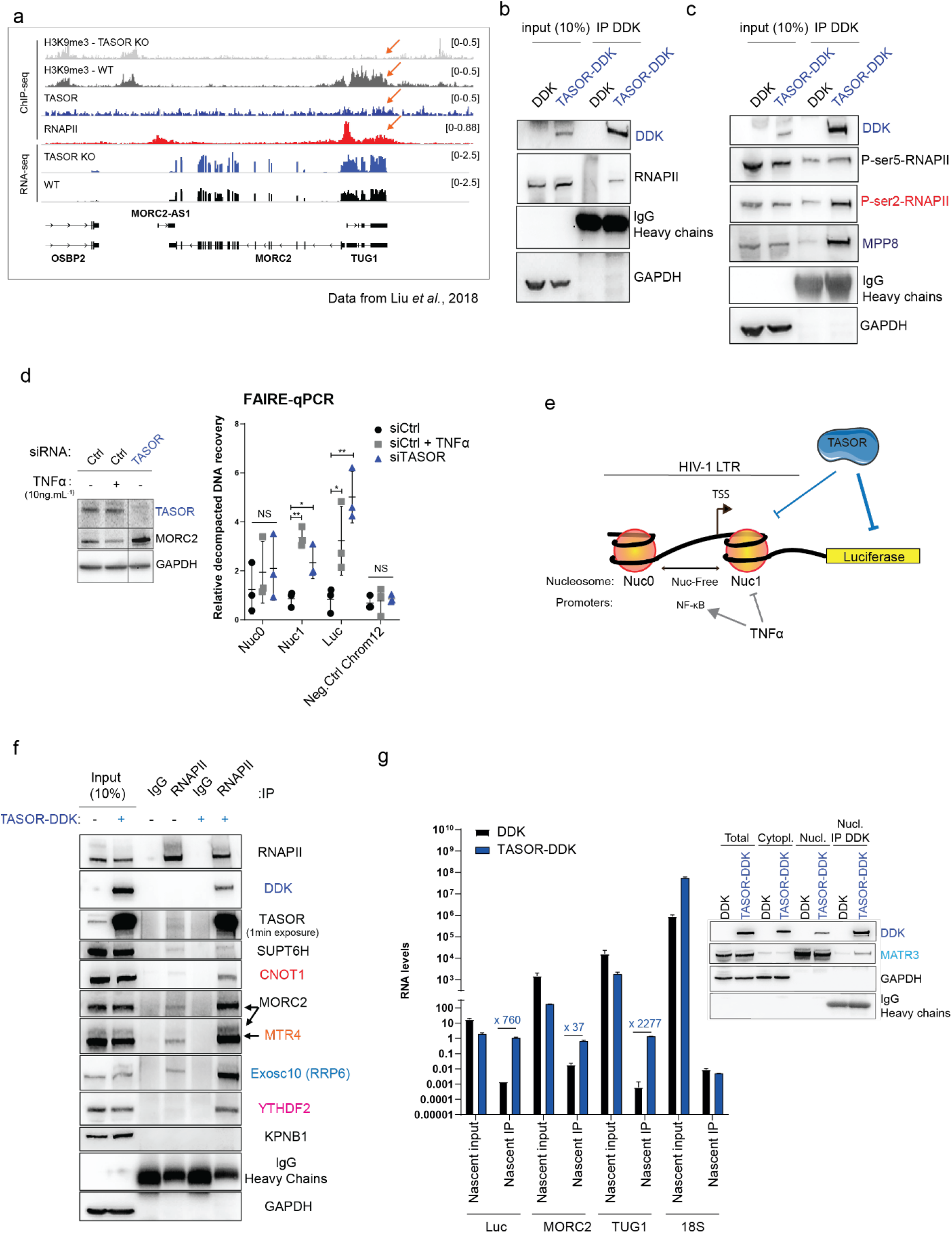
TASOR recruits RNA-degradation factors onto elongating RNAPII to silence gene expression. **a** TASOR colocalizes with HUSH-dependent H3K9me3 deposit and RNAPII on TUG1 lncRNA gene body in K562 cells. ChIPseq, RNAseq performed by Liu *et al*.,^2^ and Deposited at GEO under the accession number GSE95374 were loaded on Integrative Genomics Viewer (IGV). Orange arrow shows the colocalization signals between HUSH-dependent H3K9me3 marks, TASOR and RNAPII. **b** TASOR interacts with RNAPII. DDK, TASOR-DDK vectors were transfected in HeLa HIV-1 LTR-ΔTAR-Luc cells. After 48 h, lysates were treated and anti-DDK immunoprecipitation was performed. **c** TASOR interacts with elongating RNAPII. DDK, TASOR-DDK vectors were transfected in HeLa HIV-1 LTR-ΔTAR-Luc cells. After 48 h, anti-DDK immunoprecipitation was performed. Ser5-phosphorylated, Ser2-phosphorylated RNAPII are markers of RNAPII in the initiating and elongating phases of transcription respectively, MPP8 is a positive control. **d** TASOR depletion triggers the decompaction of the coding sequence from the HIV-1 LTR Transcription Start Site (TSS). HeLa HIV-1 LTR-Luc cells were transfected with siCtrl or siTASOR for 72h. TNFα (10ng/mL-4h) treatment that increases transcription from the TSS and then favors Nuc1 eviction was performed in siCtrl samples as a control. FAIRE-qPCR was performed (n=3), two-sided unpaired t test was applied (*P < 0.05; **P < 0.01). **e**, Schematic representation of HIV-1 LTR-Luc regions compacted by TASOR. **f** TASOR recruits CNOT1 and its partners MTR4, Exosc10, YTHDF2 onto RNAPII. DDK, TASOR-DDK vectors were transfected in HeLa HIV-1 LTR-ΔTAR-Luc cells. After 48 h, anti-RNAPII immunoprecipitation was performed. TASOR was revealed either with an anti-DDK (TASOR-DDK), or with an anti-TASOR to detect the endogenous protein as well. SUPT6H is a control of a known-RNAPII partner. MORC2 is necessary for HUSH-mediated gene silencing according to^15^. **g** TASOR binds the nascent RNA from the HIV-1 LTR and of its gene targets. DDK, TASOR-DDK vectors were transfected in HeLa HIV-1 LTR-ΔTAR-Luc cells. Nascent RNA was labelled and an anti-DDK Immunoprecipitation was performed from nuclear extract. The western-blot shows the purity of each fraction and of the immunoprecipitation. GAPDH and TASOR’s partner MATR3 are markers of the cytoplasmic and nuclear fractions respectively. Input RNAs and DDK/TASOR-DDK-bound nascent RNAs were purified, quantified by RT-qPCR. Immunoprecipitated RNA was normalized on input signal. Shown values are fold enrichment in comparison to the DDK condition.

Overall, our results support a model in which TASOR, in interaction with CNOT1, provides a platform along transcription to destabilize nascent transcripts. This conclusion is supported by (i) the effect of TASOR beyond transcription *per se* at a post-transcriptional level, (ii) the interaction of TASOR with CNOT1 and their synergistic repressive effect on HIV-1 LTR expression, (iii) the interaction and cooperation of TASOR with MTR4 and Exosc10, (iv) the fact that these interactions are stabilized in the presence of RNA, (iv) the decompaction of the coding sequence under TASOR depletion, (v) the interaction of TASOR with RNAPII, mostly under its elongating form, (vi) the recruitment of the RNA degradation factors on RNAPII when TASOR is overexpressed, and (vii) the interaction of TASOR with nascent transcripts. Keeping in mind the role of TASOR at the epigenetic level^1^, we propose a mechanism of feedback control in which TASOR and CNOT1 would follow RNAPII along transcription by association with the nascent transcript and recruit RNA metabolism proteins, leading in turn to the degradation of the transcript and the deposit of H3K9me3 marks (Fig. S5). This model is constructed by analogy to the gene expression repression models reported for the RITS complex in fission yeast or the piRNA-guided transcriptional silencing Piwi-Asterix-Panoramix-Eggless complex in drosophila^39–41^. In each case, transcriptional epigenetic repression is coupled to the synthesis of a transcript and sometimes to its degradation. The fission yeast RITS complex seems to match well our HUSH model, at least when involved in the RNAi-independent pathway, with yeast Chp1 and Tas3 sharing structural features with TASOR, MPP8 and periphilin, as also noticed recently by Schalch *et al* and Douse *et al*^37,42^ and with its ability to recruit CNOT1/NOT1^43^, the exosome and MTR4 alike TASOR. In agreement with our data, HUSH targets were shown to be enriched in transcriptionally active chromatin^2,4,42^.

Nonetheless, TASOR and CNOT1 did not seem to cooperate to repress the expression of the MORC2 gene, a cellular target of TASOR. Thus, it remains to be determined whether our conclusion using a simplified HIV transcription model holds true for the different HUSH targets alike endogenous repressed genes or exogenous targets including the murine leukemia virus that is repressed before integration by HUSH-NP220^44^. We also could not exclude different mechanisms of HUSH-mediated repression on different targets, even with the involvement of an RNAi-dependent pathway in some cases. Regarding HIV, as HUSH appears to operate along with RNA synthesis and turnover, we could suspect HUSH to be involved in the establishment of latency, when viral RNA is still produced. Since HUSH repression is dependent on the integration site^1^, it might be possible that HUSH spreads the H3K9me3 marks on the provirus sequence that has integrated in poorly, but still, transcribing regions with signals of heterochromatinization. Then HUSH could intervene by counteracting stochastic bursts of expression from the lowly expressed or latent promoter^45^. Tackling the question of HUSH role in HIV replication is going to be the next challenge, not only because of the need to find the spatiotemporal window of HUSH restriction but also because of the existence of multiple spliced and un-spliced HIV transcripts. A comprehensive view of HUSH-mediated mechanism of HIV repression might help to propose strategies for eliminating the HIV reservoir.

## Methods

### Plasmids

TASOR expression vectors pLenti-myc-DDK, pLenti-TASOR-myc-DDK were purchased from Origene. pLenti-TASOR-myc-DDK expresses a TASOR isoform of 1512 aminoacids (NCBI Reference Sequence: NP_001106207.1). DDK-CNOT7 plasmid is a kind gift from Nancy Standart. The R42A mutant of Vpx HIV-2 Ghana was produced by site-directed mutagenesis using the pAS1B-HA-Vpx HIV-2 Ghana construct as template.

### Cell lines

Cell lines were regularly tested for mycoplasma contamination: contaminated cells were discarded to perform experiments. Cells were cultured in media from ThermoFisher: DMEM (HeLa, 293T) containing 10% heat-inactivated fetal bovine serum (FBS, Dominique Dutscher), 1,000 units.ml–1 penicillin, 1,000μg.ml–1 streptomycin. HeLa LTR-ΔTAR-Luc cells were generated in the laboratory of Stephane Emiliani from the HeLa LTR-Luc cells described by du Chené *et al*., 2007^46^.

### siRNA treatment

siRNA transfections were performed with DharmaFECT1 (Dharmacon, GE Lifesciences). The final concentration for all siRNA was 100nM. The following siRNAs were purchased from Sigma Aldrich: siTASOR: SASI_Hs02_00325516; siCNOT1: SASI_Hs02_00349201; siCNOT7: SASI_Hs02_00344676; siDICER: SASI_Hs01_00160748 siMATR3: SASI_Hs02_00352277; siMTR4: SASI_Hs01_00072261 siYTHDF2 SASI_Hs01_00133218; siExosc10: SASI_Hs01_00183400. The non-targeting control siRNAs (MISSION siRNA Universal Negative Control #1, SIC001) were purchased from Sigma Aldrich.

### Luciferase activity assay

Cells were washed twice with PBS then lysed directly in wells using 1× cell culture lysis reagent (Promega). Cell lysates were clarified by centrifugation, luciferase activity was measured using a luciferase assay system (Promega) and a TECAN multimode reader Infinite F200 Pro.

### Nuclear Run On (NRO)

HeLa LTR-ΔTAR-Luc cells were grown in 10cm dishes and transfected with siRNA ctrl, TASOR + ctrl, CNOT1 + ctrl and TASOR+ CNOT1 at a final concentration of 100nM each. NRO was performed as described by Roberts and colleagues^47^, except for these specific points: At step 14, RNA extractions with rDNase treatment were performed with NucleoSpin RNA, Mini kit (740955.250, Macherey-Nagel). At Step 44, Reverse transcription was performed with Maxima First Strand cDNA Synthesis Kit with dsDNase (K1672, ThermoFisher). qPCR was finally performed as described in the RT-qPCR section.

### RT-qPCR

Except for step 14 in the *Nuclear Run On* experiments, all RNA extractions and purifications were based on a classical TRI Reagent (T9424-200ML, Merck) protocol. Reverse transcription steps were performed with Maxima First Strand cDNA Synthesis Kit with dsDNase. PCRs were performed thanks to a LightCycler480 (Roche) using a mix of 1x LightCycler 480 SYBR Green I Master (Roche) and 0.5μM primers which sequences are described in the table.

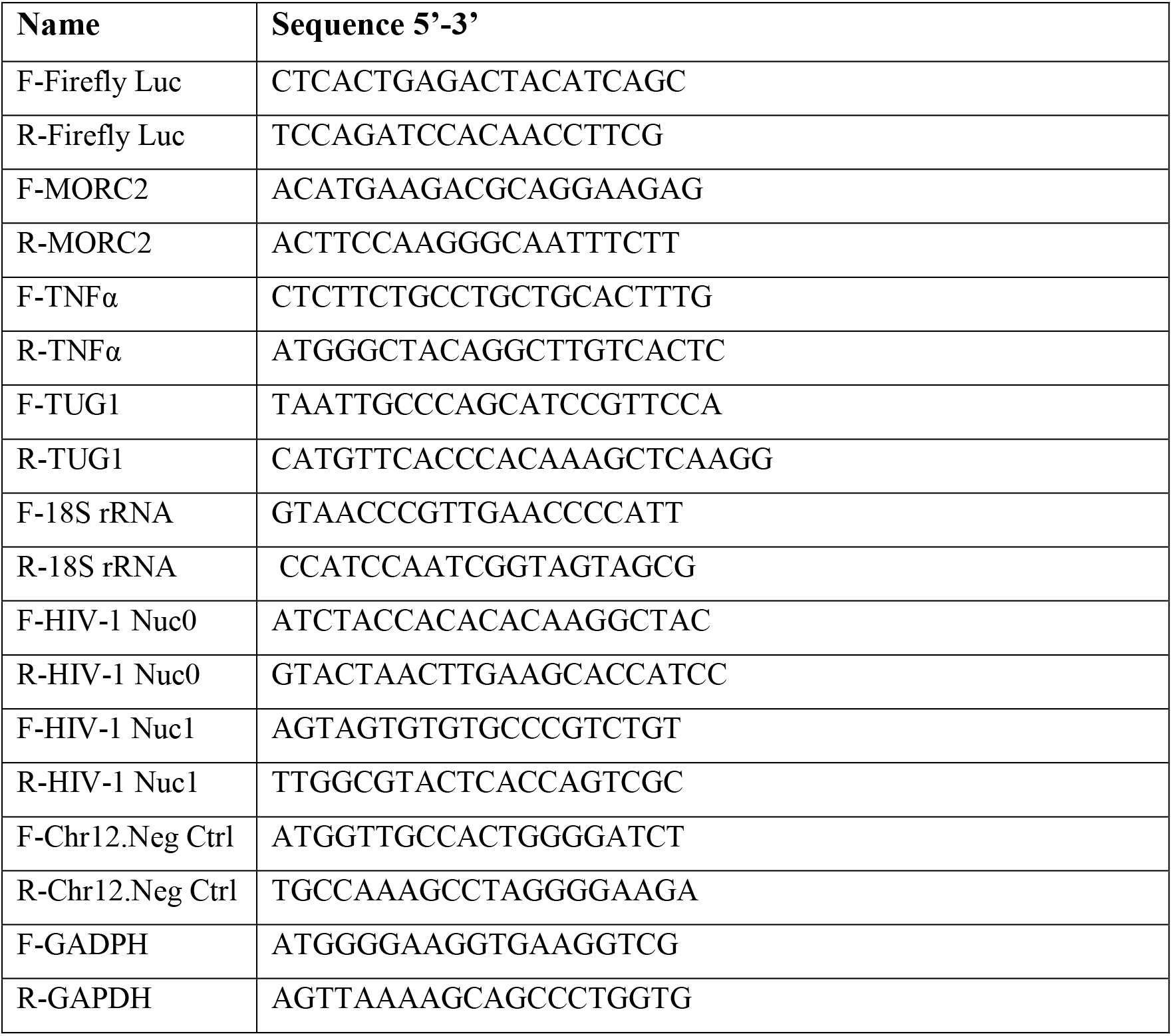

### FAIRE-qPCR

Around 6×10^6^ HeLa HIV-1 LTR-Luc cells were transfected with siRNA Ctrl or siRNA TASOR for 72h. TNFα (10 ng.mL^-1^) was added or not 4 hours before cell recovery. The FAIRE-qPCR protocol from^48^ was then followed. At step 3.2, the chromatin of each sample was sonicated with a Diagenode BIORUPTOR Pico with 10 rounds of sonication 30s on/30 s off. qPCR was performed using the primers listed in the RT-qPCR section.

### Cell fractionation

HeLa cells grown in 10cm dishes were washed with cold Dulbecco’s PBS 1x (ThermoFisher). After trypsination (25200056, ThermoFisher), cells were recovered in 1.5mL tubes and washed once with ice-cold PBS. After 4 minutes of centrifugation at 400g, 500μL of Cytoplasmic Lysis Buffer (10mM TRIS-HCl pH7.5, 10mM NaCl, 3mM MgCl_2_, and 0.5% IGEPAL^®^ CA-630 (I8896-100ML-Merck)) was added on the cell pellet and resuspended pellet was incubated on ice for 5min. Cells were then centrifuged at 300g for 4min at 4°C. The supernatant was saved for cytoplasmic fraction. The pellet resuspended with 1mL of Cytoplasmic Lysis Buffer and re-centrifuged at 300g for 4min at 4°C twice. Finally, the nuclear pellet was lysed with 200μL of RIPA Buffer.

### Immunoprecipitation, Western Blot procedures and Antibodies

For TASOR-DDK or DDK-CNOT7-immunoprecipitations: HeLa/293T cells grown in 10 cm dishes were transfected with pLenti-DDK or with pLenti-TASOR-DDK or Flag-CNOT7 plasmids with CaCl2. 48h after transfection, cells were lysed in 500μl RIPA buffer (50mM Tris-HCl pH7.5, 150mM NaCl, 1mM MgCl_2_, 1mM EDTA, 0.5% Triton X100) containing an anti-protease cocktail (A32965, ThermoFisher). Cell lysates were clarified by centrifugation and a minimum of 200μg of lysate was incubated with pre-washed EZview™ Red ANTI-FLAG^®^ M2 Affinity Gel (F2426, Merck) at 4°C, under overnight rotation. After three washes in wash buffer (50mM Tris-HCl pH7.5, 150mM NaCl), immunocomplexes were eluted with Laemmli buffer 1× and were separated by SDS-PAGE. For endogenous Immunoprecipitation, the same procedures were followed but a minimum of 400μg of lysate was used for overnight IP with 4μg of the corresponding antibody and pre-washed Pierce™ Protein A/G Magnetic Beads (88802, ThermoFisher) were added for 1h at room temperature. Following transfer onto PVDF membranes, proteins were revealed by immunoblot. Signals were acquired with Fusion FX (Vilber). The following antibodies, with their respective dilutions in 5% skimmed milk in PBS-Tween 0.1%, were used: anti-Flag M2 (F1804-200UG, lot SLCD3990, Merck) 1/1000; anti-TASOR (HPA006735, lots A106822, C119001, Merck) 1/1000; anti-MPP8 (HPA040035, lot R38302, Merck) 1/500; anti-CNOT1 (For WB: 66507-1-Ig, Proteintech, 1/1000; For IP: 14276-1-AP, Proteintech); anti-CNOT7 (14102-1-AP, Proteintech) 1/500; anti-CNOT9 (22503-1-AP, Proteintech) 1/500; anti-DHX9 (17721-1-AP, Proteintech) 1/1000; anti-Exosc10 (11178-1-AP, Proteintech) 1/1000; anti-HLTF (ab17984, Abcam) 1/1000; anti-IGF2BP1 (22803-1-AP, Proteintech) 1/1000; anti-KPNB1 (HPA029878-100ul, lot D114771, Merck) 1/1000; anti-MATR3 (12202-2-AP, Proteintech) 1/1000; anti-MORC2 (PA5-51172, TermoFisher) 1/1000; anti-MTR4 (For WB and IP: 12719-2-AP, Proteintech) 1/1000; anti-PPHLN-1 (HPA038902, Lot A104626, Merck) 1/1000; anti-RNAPII (For IP and WB: F12, sc-55492, Santa Cruz Biotechnology) 1/1000; anti-Ser2P-RNAPII (13499S, Cell Signaling technology) 1/1000 in 2.5% BSA-TBS-Tween 0.1%; anti-Ser5P-RNAPII (13523S, Cell Signaling technology) 1/1000 in 2.5% BSA-TBS-Tween 0.1%; anti-SUPT6H (For WB and IP: 23073-1-AP, Proteintech) 1/1000; anti-YTHDF2 (24744-1-AP, Proteintech) 1/1000; anti-Actin (AC40, A3853, Merck) 1/1000; anti-αTubulin (T9026-.2mL, lot 081M4861, Merck) 1/1000; anti-GAPDH (6C5, SC-32233, Santa Cruz) 1/1000. All secondary antibodies anti-mouse (31430, lot VF297958, ThermoFisher) and anti-rabbit (31460, lots VC297287, UK293475 ThermoFisher) were used at a 1/10000 dilution before reaction with Immobilon Forte Western HRP substrate (WBLUF0100, Merck Millipore).

### Nascent RNA IP

HeLa LTR-ΔTAR-Luc were grown in 10cm dishes and transfected with pLenti-DDK or pLenti-TASOR-DDK. 48 hours post-transfections, the cells were recovered and labeling of nascent RNAs was undertaken with the *Nuclear Run On* protocol^47^. 10% of nuclei were put aside for purification of BrUTP incorporated Nascent RNAs (input). From the remaining nuclei RNA-IP was then undertaken with the MagnaRIP Magna RIP™ RNA-Binding Protein Immunoprecipitation Kit protocol (17-700, Merck) from step I.3. The EZview™ Red ANTI-FLAG^®^ M2 Affinity Gel (F2426, Merck) used for DDK immunoprecipitation was washed with the RIP wash buffer and was incubated with Nuclei Lysates at 4°C overnight. From step III.6 of the MagnaRIP protocol, 10% of IPed lysate was recovered and washed three times with IP wash buffer (50mM Tris-HCl pH7.5, 150mM NaCl) and loaded on SDS-Page gel to check DDK pull-down efficiency. The remaining IPed Lysate was then washed from step III.6 of the MagnaRIP protocol. Finally, purification of BrUTP incorporated IPed Nascent RNAs was undertaken from step 25 of the *Nuclear Run On* protocol^47^. After purification, RT-qPCR of all purified RNAs was performed with primers listed in the RT-qPCR section.

## Acknowledgements

We thank all the Retrovirus, Infection and Latency group members at Institut Cochin for fruitful discussions. We warmly thank Nancy Standart for providing the Flag-CNOT7 plasmid. This work was supported by grants from the “Agence Nationale de la Recherche sur le SIDA et les hépatites virales” (ANRS) and SIDACTION. R.M. was supported by ANRS and SIDACTION. B.M. is supported by ENS Paris Saclay and P.L. by Université de Paris.

## Author Contributions

R.M and F.M-G conceived the study. R.M. and F.M-G designed experiments and interpreted data. S.E and S.G.M brought their expertise in regular discussions. R.M., M.M., P.L, B.M performed experiments. F.B re-analyzed previously published ChIPseq, RNAseq data. R.M and F.M-G wrote the paper.

**No competing financial interest**

---

**Figure S1:**
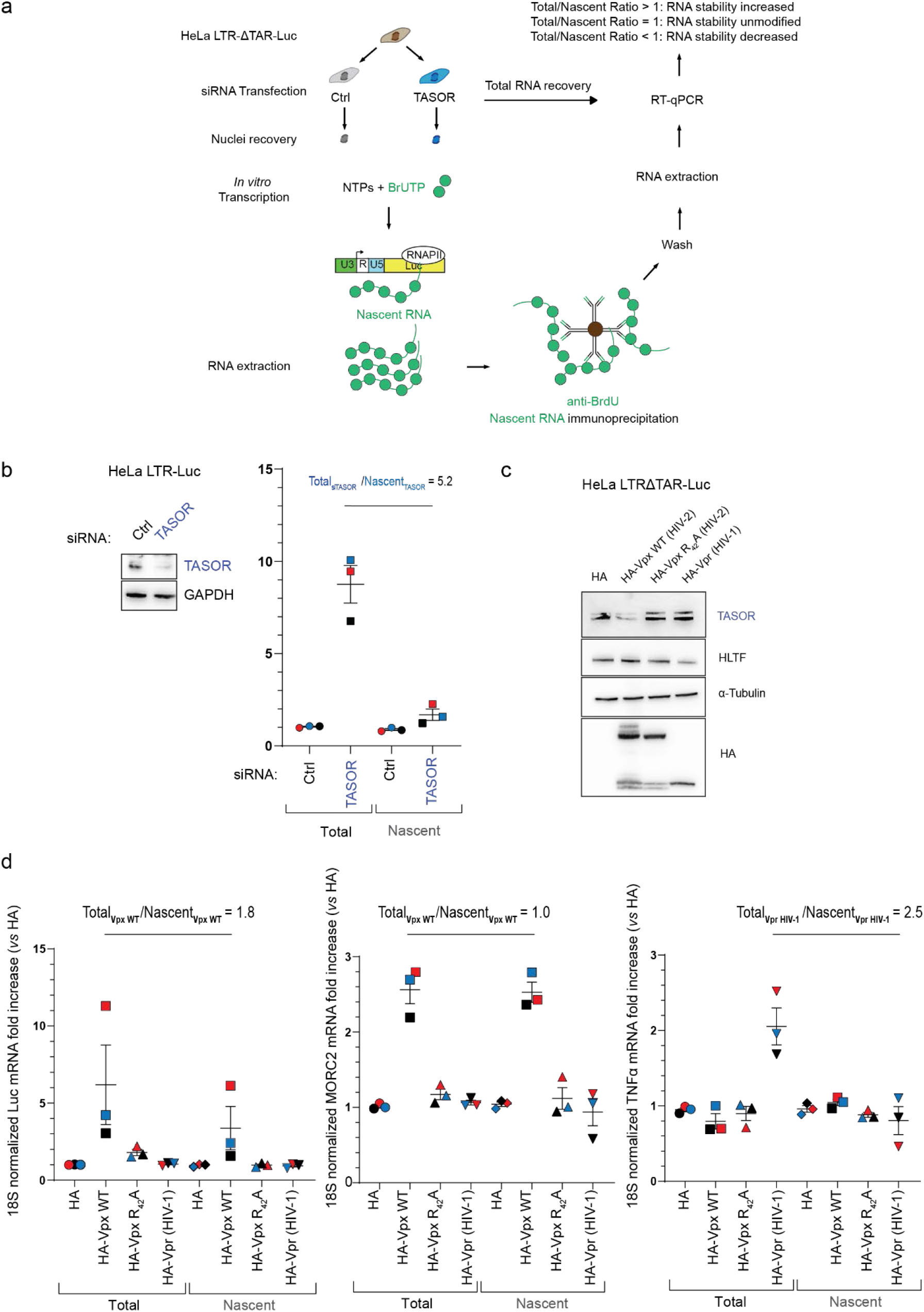
**a** *Nuclear Run On* strategy employed to characterize TASOR’s function in the mRNA metabolism pathway HeLa HIV-1 LTRΔTAR-Luc cells. **b** TASOR negatively impacts LTR-driven Luc transcript at a post-transcriptional step. *Nuclear Run On* performed in HeLa HIV-1 LTR-Luc after 72h of siCtrl or siTASOR transfection. (n=3; each color represents one different independent experiment, mean and SEM are showed). **c, d** HIV-2 Vpx mimics siRNA-mediated TASOR depletion. HeLa HIV-1 LTRΔTAR-Luc were transfected with pAS1B-HA pAS1B-HA-Vpx WT HIV-2, or pAS1B-HA-Vpx R42A HIV-2, or pAS1B-HA-Vpr HIV-1 for 48h. HLTF, target of HIV-1 Vpr^49^ is a positive control for Vpr activity. MORC2 upregulation is a control for Vpx-mediated TASOR degradation^10,15^. TASOR, HLTF, MORC2, and HA-Vpx/Vpr expressions were detected by Western-Blot (c) and *Nuclear Run On* was undertaken to measure Luc, MORC2, TNFα RNA levels at the transcriptional or post-transcriptional steps (d). (n=3; each color represents one different independent experiment, mean and SEM are showed).

**Figure S2:**
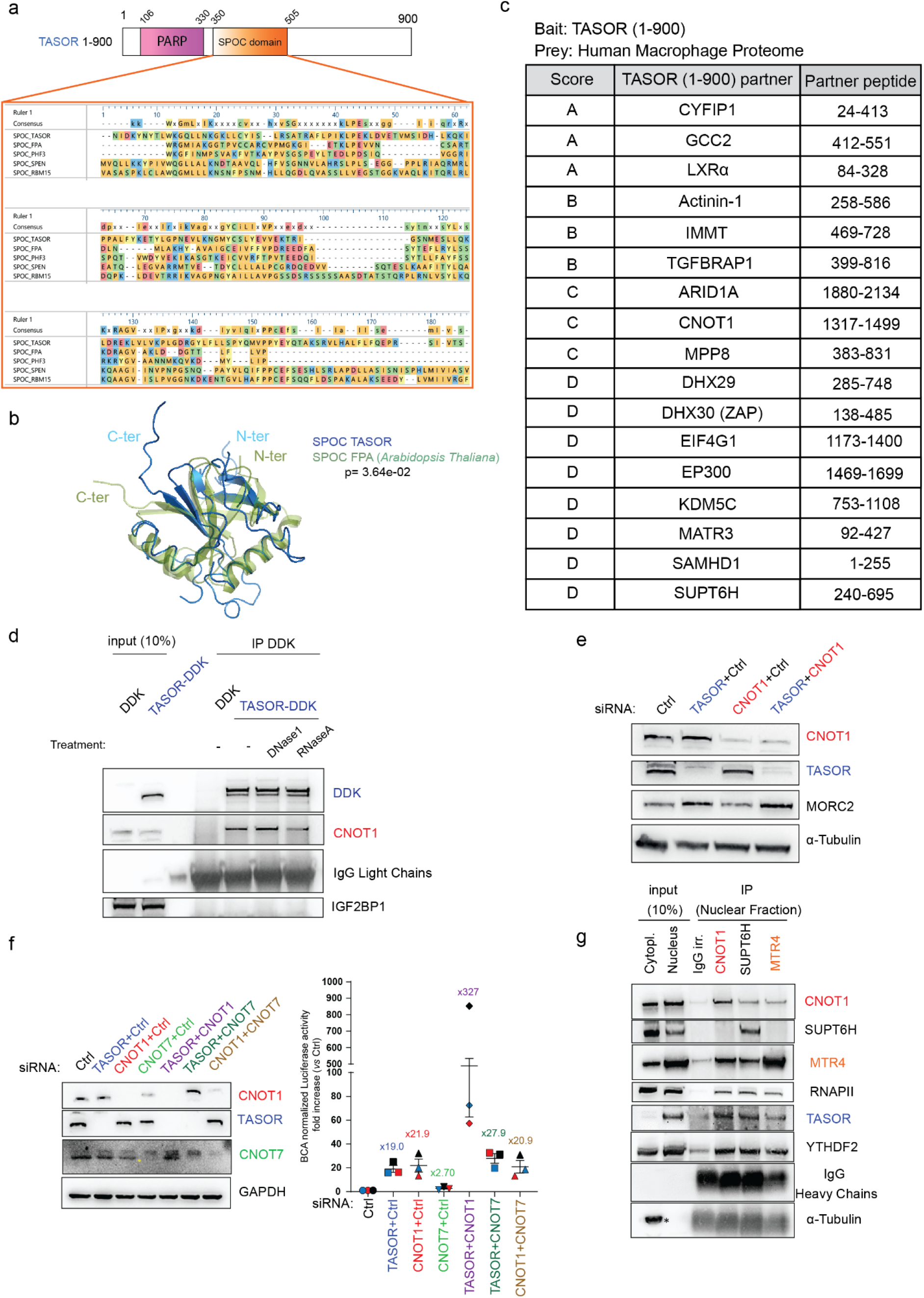
**a** Structure prediction of TASOR first 900 amino-acids detected a PARP domain and a SPOC domain. SPOC domains from different SPOC containing proteins were aligned using clustal Omega. Proteins and their identifier: *Homo sapiens* TASOR isoform1: Q9UK61-1; *Arabidopsis thaliana* FPA isoform1: Q8LPQ9-1; *Homo sapiens* PHF3 isoform1: Q92576-1; Homo sapiens SPEN: Q96T58-1; *Homo sapiens* RBM15 isoform1: Q96T37-1. **b** Overlay of TASOR SPOC domain predicted structure (RaptorX) with the *Arabidopsis thaliana* FPA SPOC domain (PDB:5KXF). **c** A yeast two-hybrid (Y-2-H) screen was performed using a human macrophage cDNA library to identify interacting proteins with TASOR (1-900). Interacting partners were assigned a predicted biological score from A-F to assess the confidence of an interaction being specific (with A indicating very high confidence, and F indicating experimentally determined artifacts). TASOR’s interaction site (aa) on the identified partner is shown in the ‘partner peptide’ column. **d** The interaction between TASOR and CNOT1 is favored in presence of a transcript. The same experiment as presented in Fig. 2c was conducted in HeLa HIV-1 LTRΔTAR-Luc cells. IGF2BP1 is a negative control. **e** Western-blot associated with the *Nuclear Run On* data presented on Fig. 2. **f** Cooperation between the CCR4-NOT complex deadenylase CNOT7 and TASOR in the repression of HIV-1 LTR-Luc expression. Following 72h of siRNA transfections in HeLa HIV-1 LTR-ΔTAR-Luc, cells were lysed, siRNA-mediated depletion of proteins was controlled by Western-Blot and Luciferase activity was measured and normalized on protein concentration. (n=3; each color represents one different independent experiment, mean and SEM are showed). **g** Endogenous TASOR interacts with the histone chaperone and transcription elongation factor SUPT6H. HeLa HIV-1 LTRΔTAR-Luc cells were fractionated and CNOT1, SUPT6H, MTR4 immunoprecipitations were performed from nuclear extracts.

**Figure S3:**
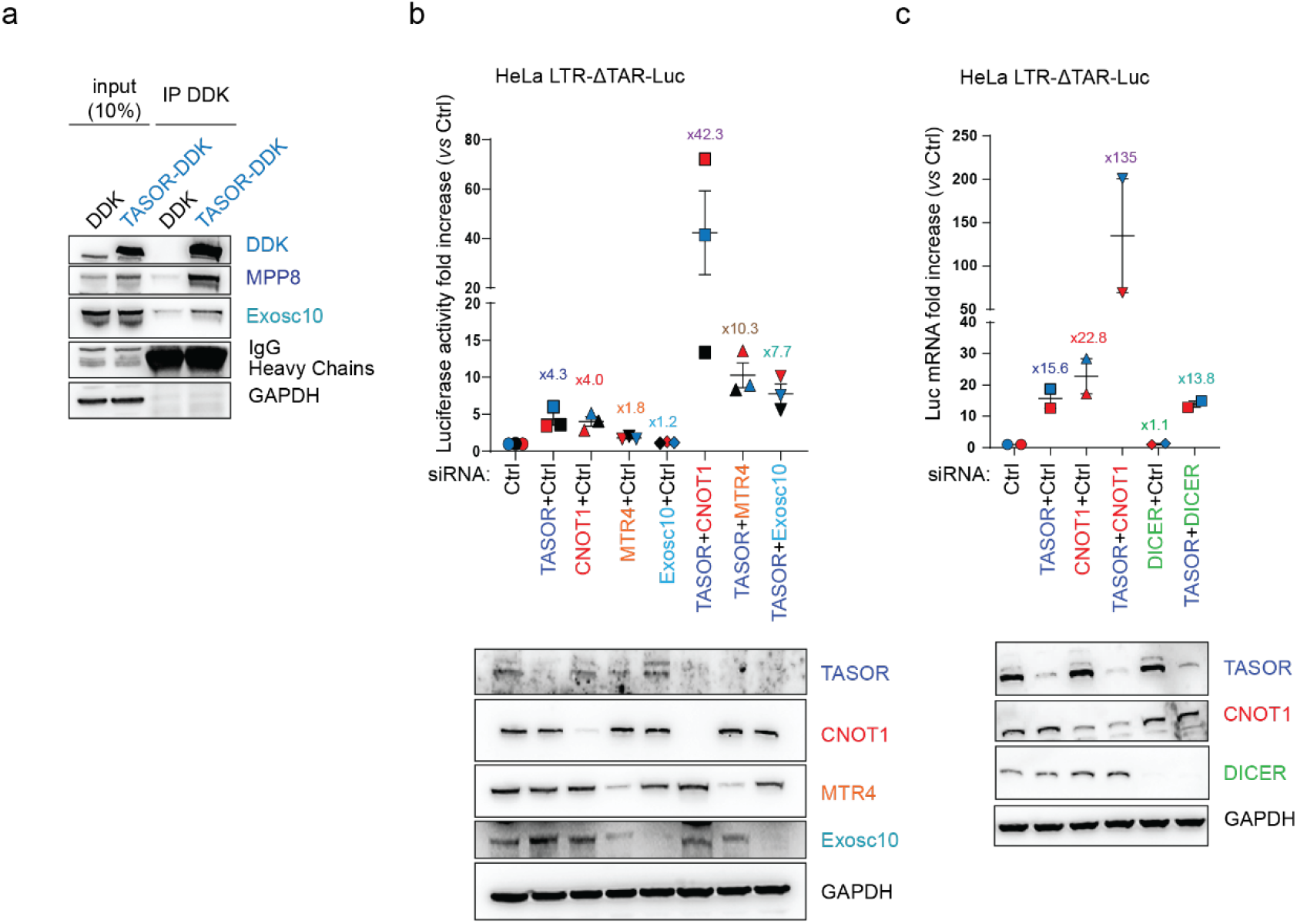
**a** TASOR interacts with the exosome factor Exosc10. HeLa HIV-1 LTRΔTAR-Luc cells were transfected with DDK or TASOR-DDK constructs for 48h and a DDK immunoprecipitation was performed. **b** TASOR cooperates with TRAMP-like/NEXT components MTR4 and Exosc10 in the repression of HIV-1 LTR-Luc expression. After 72h of siRNA transfections in HeLa HIV-1 LTR-ΔTAR-Luc, cells were lysed, siRNA-mediated depletion of proteins was controlled by Western-Blot and Luciferase activity was measured and normalized on protein concentration. (n=3; each color represents one different independent experiment, mean and SEM are showed). **c** RNAi-pathway factor DICER has no role and does not cooperate with TASOR in the repression of the HIV-1 LTR-Luc expression. After 72h of siRNA transfections in HeLa HIV-1 LTR-ΔTAR-Luc, cells were lysed, siRNA-mediated depletion of proteins was controlled by Western-Blot and Luciferase activity was measured and normalized on protein concentration. (n=2; each color represents one different independent experiment, mean and SEM are showed).

**Figure S4:**
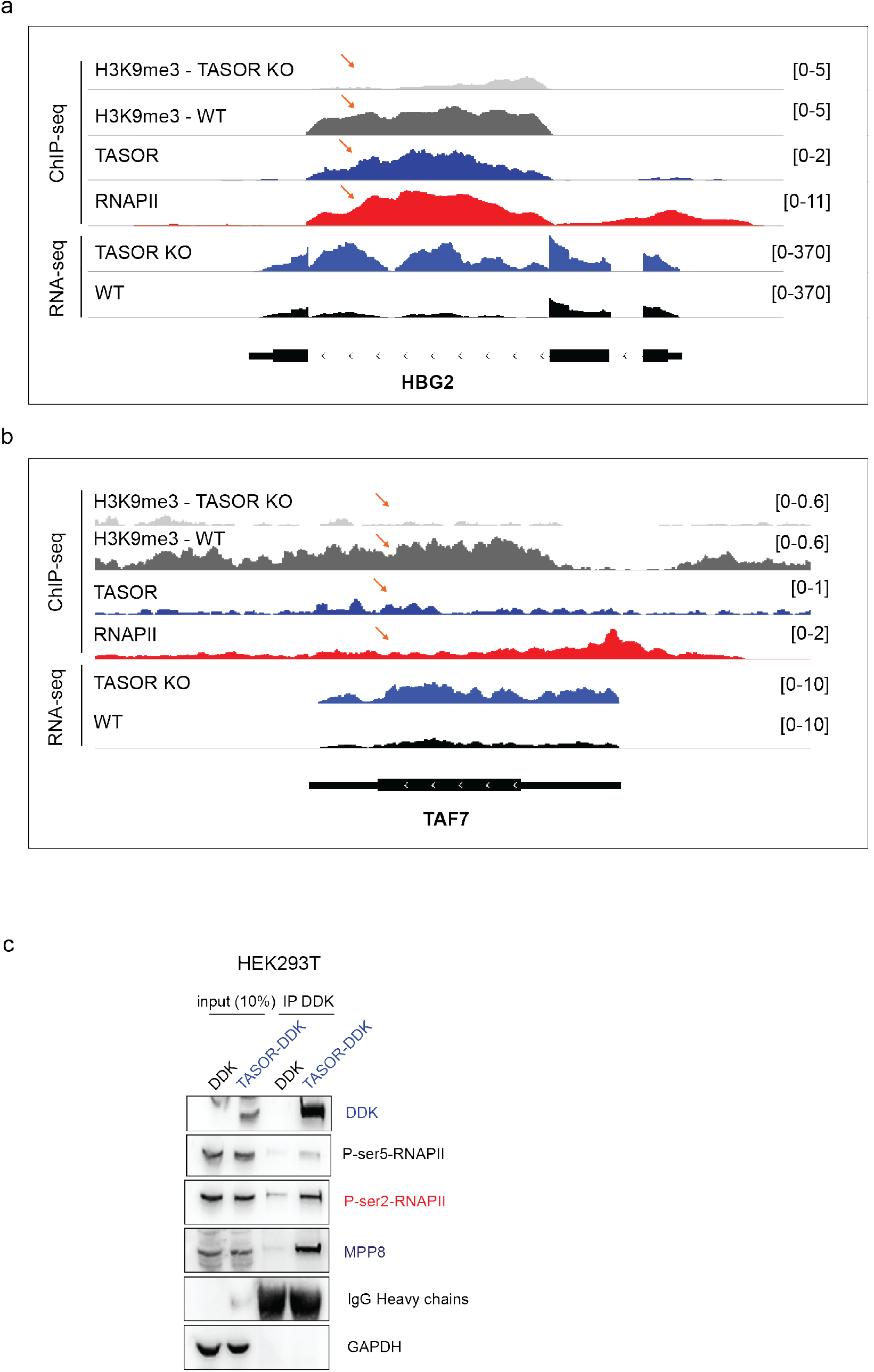
**a** TASOR colocalizes with HUSH-dependent H3K9me3 deposit and RNAPII on the intron of HBG2 gene in K562 cells. ChIPseq, RNAseq performed by Liu *et al*.,^2^ and Deposited at GEO under the accession number GSE95374 were loaded on Integrative Genomics Viewer (IGV). Orange arrow shows the colocalization signals between HUSH-dependent H3K9me3 marks, TASOR and RNAPII. **b** TASOR colocalizes with HUSH-dependent H3K9me3 deposit and RNAPII on the TAF7 gene body in K562 cells. ChIPseq, RNAseq performed by Liu *et al*.,^2^ and Deposited at GEO under the accession number GSE95374 were loaded on Integrative Genomics Viewer (IGV). Orange arrow shows the colocalization signals between HUSH-dependent H3K9me3 marks, TASOR and RNAPII. **c** TASOR interacts with elongating RNAPII. DDK, TASOR-DDK vectors were transfected in HEK293T cells. After 48 h, anti-DDK immunoprecipitation was performed. Ser5-phosphorylated, Ser2-phosphorylated RNAPII are markers of RNAPII in the initiating and elongating phases of transcription respectively, MPP8 is a positive control.

**Figure S5:**
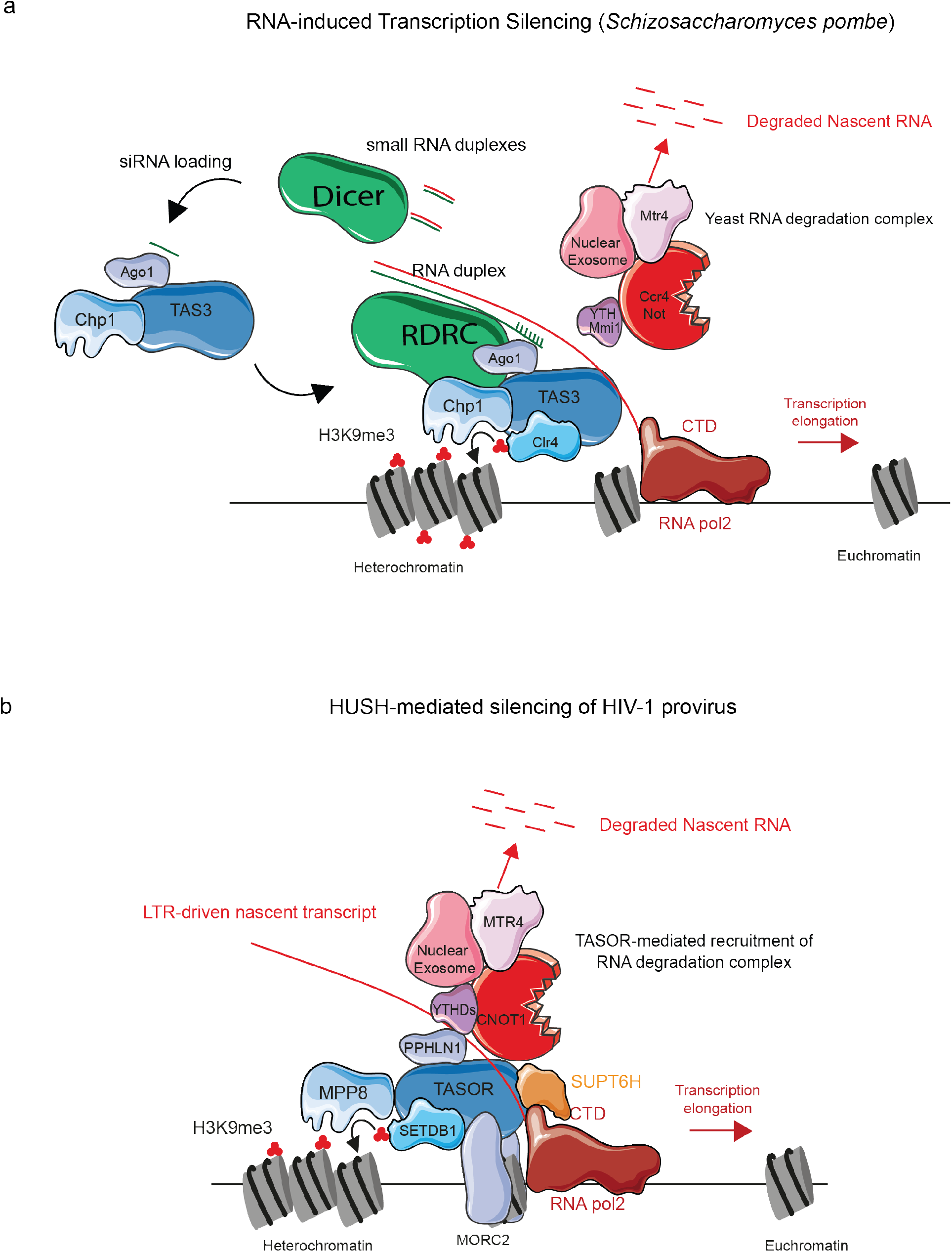
Overall conserved model of retroelement repression by Fission Yeast RITS complex and Human HUSH complex. **a** Retroelement silencing in *Schizosaccharomyces pombe:* Chp1, Tas3, Ago1 build the RITS complex which, with the help of the Histone methyl transferase Clr4, mediates retroelement silencing by spreading the H3K9me3 marks in an RNAi-dependent pathway (RDRC/DICER/siRNA) and associates with Mmi1, Mtr4, nuclear exosome in an RNAi-independent silencing pathway. **b** Integrated HIV-1 LTR provirus silencing in *Homo sapiens*: The HUSH complex interacts and follows elongating RNAPII, recruits RNA degradation factors to degrade the LTR-driven Luc RNA (this study) and spreads the deposit of H3K9me3 on the coding sequence as already shown in^3^.

